# Transitions in sex determination mechanisms through parental antagonism

**DOI:** 10.1101/2023.08.04.551945

**Authors:** Martijn A. Schenkel

**Affiliations:** Department of Biology, Georgetown University, 37th and O St. NW, Washington DC, 20057, USA; Groningen Institute of Evolutionary Life Sciences, University of Groningen, Nijenborgh 7, 9747 AG, Groningen, The Netherlands

## Abstract

Parental antagonism (PA) occurs when the fitness effects of a gene depend on the parent from which it is inherited. Such genes may become enriched on sex chromosomes, due to their biased inheritance patterns. Although various sex determination (SD) genes exhibit parent-of-origin effects themselves, and between-parent conflict over offspring sex may affect SD, PA itself has not been considered as a driver of SD transitions. Here, I present a model to investigate the scope for transitions in SD mechanisms through PA. My model assumes an ancestral SD locus linked to a PA gene, as well as an autosomal PA gene in whose vicinity a novel SD gene arises. Transitions between functionally-homologous genes are found to depend on the fitness effects of both PA genes and their linkage to nearby SD genes. Transitions between male and female heterogamety by the invasion of a dominant SD gene are however nearly unconstrained. This also allows for back-and-forth dynamics where the ancestral SD and novel SD genes constantly evolve to be dominant over each other. These results further underline the malleability of SD mechanisms, and the need to consider parent-of-origin effects in driving transitions in SD, through proximate and/or ultimate means.

## Introduction

Sex determination directs the development of an individual into a female or male, and its proper execution is therefore essential to ensure the developing individual will be able to reproduce. Despite this pivotal role, the mechanisms controlling sex determination are liable to turnover. Such transitions in sex determination may be driven by neutral processes (Bull & Charnov, 1977; Veller *et al*., 2017) as well as a variety of selective processes such as segregation distortion, sex ratio selection, and sexually antagonistic selection (reviewed in van Doorn 2014). Furthermore, environmental effects may impinge on sex determination, and thereby help shape its evolutionary trajectory (e.g. Pen et al. 2010; Schenkel et al. 2023). Of particular interest here, parent-offspring conflict as well as between-parent conflict over the offspring sex (or sex ratio) can profoundly affect sex determination as well (e.g. Pen 2006; Uller et al. 2007). Proximate studies of genetic sex determination cascades has furthermore revealed that sex determination processes involve parent-of-origin effects, such as the imprinting mechanism that controls the activity of sex determination genes in in the parasitoid wasp *Nasonia vitripennis* (Verhulst *et al*., 2013; Zou *et al*., 2020), and maternal provisioning of *tra* mRNA that kickstarts feminization in the housefly *Musca domestica* (Dübendorfer & Hediger, 1998; Hediger *et al*., 2010). Nonetheless, the possibility for genes that exhibit parent-of-origin effects on fitness have not yet been implicated as a driver of sex determination transitions.

Sex chromosomes occupy a special niche within the genome owing to the presence of the master sex determination gene, segregating to and from females and males in biased patterns (Haig et al. 2014; Schenkel and Beukeboom 2016; Figure 1A). Consequently, genes that have different effects on fitness depending on the sex of the carrier, or the origin of the allele, may evolve differently on these chromosomes as compared to when they occur on autosomes (Patten & Haig, 2009; Jordan & Charlesworth, 2012). Genes that differently affect fitness in females and males are known as sexually antagonistic genes, or are said to be involved in intralocus sexual conflict (Schenkel *et al*., 2018); similarly, genes with different effects depending on their parent of origin are known as parentally antagonistic genes (Haig, 1997). Sexually antagonistic genes have previously been shown to be able to drive transitions in sex determination, i.e. the invasion of a novel sex determination gene, which in doing so turns a former autosome into a novel sex chromosome pair (van Doorn & Kirkpatrick, 2007, 2010; van Doorn, 2014). Here, selection may favor the evolution of linkage disequilibrium between a gene with male-beneficial/female-detrimental effects to an allele that causes maleness. Such a supergene would more often end up in males, in whom it enhances fitness, and less often in females, in which it exerts a fitness cost. Selection similarly favors its counterpart, which consists of a recessive female-determining allele and an allele with female-beneficial/male-detrimental effects. Possibly, parentally antagonistic selection may favor similar supergenes as a gene that causes maleness, and hence is inherited paternally, becomes associated with an allele that has fitness benefits when inherited paternally; inversely, the recessive feminizing allele would become associated with genes that confer fitness benefits when maternally inherited.

**Figure 1:**
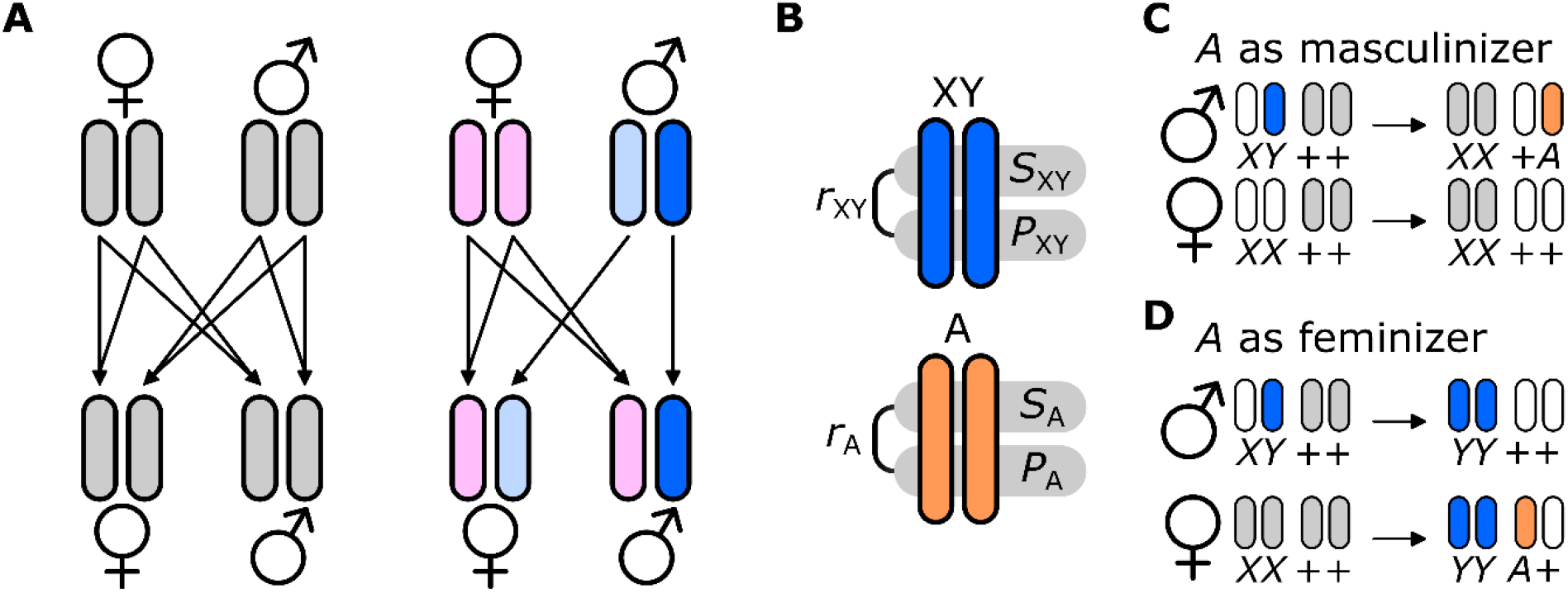
Sex chromosome inheritance, model setup, and transitions in sex determination. (A) Autosomal inheritance (left) allows for all chromosomes to freely segregate from and to females and males (indicated by arrows). Sex chromosomal inheritance (right) leads to biased patterns: the Y-chromosome (dark blue) is always transmitted paternally from fathers to sons; the X-chromosome in such fathers (light blue) is always transmitted to daughters. X-chromosomes in mothers (pink) are transmitted to daughters and sons alike. (B) In my model, I consider two chromosome pairs XY (blue) and A (orange), each of which carries a (potential) sex determination locus *S* and a parentally antagonistic locus *P*; recombination occurs at a rate r between them. (C) If *A* has a male-determining function, transitions in sex determination occur via the loss of *Y* (and fixation of *X*) as *A* invades on the paternal copy in males. (D) If *A* has a female-determining function, transitions in sex determination occur via the fixation of *Y* in both sexes, and the invasion of *A* on the maternal copy in females. Grey chromosomes indicate autosomes; white chromosomes indicate sex chromosome complements that lack a dominant sex-determining allele *Y* or *A*.

Here, I present a model of transitions in sex determination through linkage between parentally antagonistic genes and sex determination genes. In line with the results from van Doorn and Kirkpatrick (2007, 2010) for transitions mediated through sexually antagonistic selection, I hypothesized that the scope for invasion of a novel sex determination gene would be positively affected by the strength of selection acting on both parentally antagonistic loci, as well as the degree of linkage between the sex determination and parentally antagonistic loci. The effect of selection here depends on the selective effect of the parentally antagonistic locus in homozygotes, as well as the different scaling parameters that determine the relative fitness of the heterozygotes in which the focal allele is maternally versus paternally inherited. I additionally consider the role of varying degrees of linkage between sex-determining genes and paternally-antagonistic genes in shaping the scope for turnover.

## Methods

### Model overview

I provide here a general description of the model; the mathematical model is explained in detail in the Supplementary Material; an overview of all model parameters including standardized values is included in Supplementary Table 1. The model is a modified version of the one presented in Schenkel et al. (2021). I present a two-locus, two-allele model per linkage group, with a genome consisting of two linkage groups (XY and A) so that the full model features four loci (Figure 1B). All individuals are diploid; all genotypes in the model are represented with the maternally-inherited first, and the paternally-inherited allele second. Each linkage group *i* carries an SD gene *S*_*i*_ and a parentally antagonistic locus *P*_*i*_ with alleles *p*_*i*1_ and *p*_*i*2_. Recombination between *S*_*i*_ and *P*_*i*_ occurs at a rate *r*_*i*_ in both sexes. The locus *S*_*XY*_ carries two alleles *X* and *Y*, and is the ancestral SD locus; the SD system is assumed to be male heterogametic (females *XX*, males *XY*). The locus *S*_*A*_ is initially fixed for an allele *+* that does not affect sex; SD transitions occur by the invasion of a novel SD allele on *S*_*A*_, which I denote *A*. If *A* has a male-determining effect, its invasion would cause an SD transition between two different male heterogamety systems (Figure 1C, Supplementary Figure 1A); females would retain an *XX; ++* genotype, but males would be *XX; +A*. If *A* has a female-determining effect, I assume it to be dominant over *Y*. Under these conditions, its invasion would lead to fixation of *Y* in all individuals and *A* on the maternal copy in females (females *YY; A+*; males *YY; ++*; Figure 1D, Supplementary Figure 1B). Note that terms such as ‘female’ and ‘male’, and similarly ‘maternal’ and ‘paternal’ can be swapped so that different SD transitions are accounted for in my model (e.g., one might label *Y* a female-determiner, so that the original SD system is female heterogametic instead of male heterogametic (females *YX*, males *XX*)).

### Parentally-antagonistic selection

Alleles *p*_*i*1_ are beneficial when maternally inherited, but detrimental when paternally inherited and vice versa for *p*_*i*2_, so that the optimal genotype for each locus *P*_*i*_ is *p*_*i*1_*p*_*i*2_ whereas the least optimal genotype is *p*_*i*2_*p*_*i*1_. The fitnesses *w*_*ijk*_ of a genotype *p*_*ij*_ *p*_*ik*_ are determined as follows:

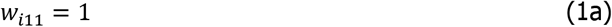

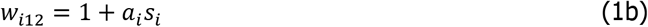

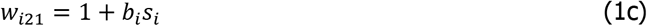

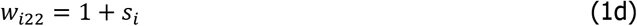

Here, *s*_*i*_ indicates the fitness of effect the *p*_*i*2_ allele, and *a*_*i*_ and *b*_*i*_ represent scaling parameters for heterozygous genotypes when *p*_*i*2_ is maternally (*a*_*i*_) or paternally inherited (*b*_*i*_). Because I assume that *p*_*i*2_ is detrimental when maternally-inherited but beneficial when paternally-inherited, this means that *a*_*i*_*s*_*i*_ < 0 and *b*_*i*_*s*_*i*_ > *s*_*i*_. For simplicity, I assume selection on *p*_*i*2_ is positive (i.e. *p*_*i*2_*p*_*i*2_ has a higher fitness than *p*_*i*1_*p*_*i*1_), so that *s*_*i*_ > 0 and therefore *a*_*i*_ < 0 and *b*_*i*_ > 1. Fitness effects of loci *P*_*XY*_ and *P*_*A*_ are multiplicative, i.e. *w* = *w*_*XY*_ × *w*_*A*_. Selection takes place during maturation from the juvenile stage into the adult stage, so that survival is proportional to relative fitness.

### Simulation procedure

I initialize a population with equal sex ratios, in which all females have a genotype *XX; ++* and males have *XY; ++* for the two (potential) sex-determining loci. Both *P* loci start with an initial frequency *p* = 0.5, regardless of the parameter values for *a*_*i*_, *b*_*i*_, *s*_*i*_, and *r*_*i*_. The values of these parameters were varied in different sets of simulations in which some were kept at constant values and others were varied by sampling from uniform distributions (for details, see results).

I restrict my analysis to cases where both parentally antagonistic loci would remain polymorphic when autosomal, i.e. both alleles *p*_*i*1_ and *p*_*i*2_ have a non-zero frequency even when not linked to a sex-determining gene. This occurs when (Pearce & Spencer, 1992; Úbeda & Haig, 2004):

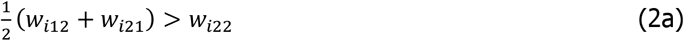

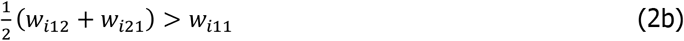

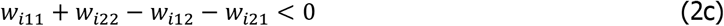

In the configuration presented here and assuming *s*_*i*_ > 0, inequalities 2a-c are satisfied when:

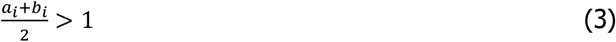

I standardize *b*_*i*_ = −*c*_*i*_*a*_*i*_ + 2 with *c*_*i*_ > 1 in each set of simulations, so that the value of *b*_*i*_ follows from the value of *a*_*i*_ and inequality 3 is satisfied.

I perform different sets of simulations where I vary the values of *a*_*i*_, *c*_*i*_, *s*_*i*_, and *r*_*i*_ in different combinations to explore how they affect the scope for invasion (see Results for details). I allow both *P* loci to evolve towards their equilibrium state during 10,000 generations, after which I introduce the *A* allele by mutating a small proportion (*p* = 0.001) of the *S*_A_ loci from *+* to *A*, proportionally distributed among all haplotypes present at non-zero frequencies in the gamete pool to prevent linkage disequilibrium. Subsequently, I allow for at least 40,000 more generations before I determine the final frequency of the *A* allele and thereby whether an SD transition has occurred. I consider scenarios where *A* represents (1) a male-determining gene that functions similarly to *Y* so that invasion of *A* entails a transition between homologous male heterogametic systems, and (2) a female-determining gene that is dominant over *Y* so that invasion of *A* entails a transition from male heterogamety to female heterogamety.

### Statistical analysis

All simulations, data analyses, and data visualization were performed in R (v. 4.2.1; (R Development Core Team, 2023)) and RStudio (v. 2023.06.01; (RStudio Team, 2023)) using the ‘tidyverse’ (Wickham *et al*., 2019), ‘mgcv’ (Wood, 2017), and ‘viridis’ (Garnier, 2018) packages. Allele frequencies for *Y* and *A* on maternal c.q. paternal copies typically evolve to values very close (but not equal) to 0 or 1; to facilitate analysis, I round these frequencies to the nearest integer to convert these to binomially-distributed data. Next, I fitted generalized additive models with binomial distributions to the rounded frequencies to interpolate between sampled parameter values. For transitions with *A* as a male-determiner, I used the frequencies of *A* on the paternal copy in males, whereas if *A* functioned as a female-determiner, I used the frequencies on the maternal copy in females because only for these copies can *A* achieve a non-zero frequency under these conditions. I used full tensor smooths between different parameter combinations as the predictor variable, depending on which parameters were varied in a set of simulations (details included with results). Thin plate regression splines with extra shrinkage were used as the base functions.

## Results

### Transitions between different male heterogamety systems

When *A* represents a male-determining gene that is functionally homologous to *Y*, its invasion means that the ancestral male heterogamety system is replaced by a similar male heterogamety system (Figures 1C, Supplementary Figure 1A). I find that the scope for such transitions depends both on the selection regimes acting on *P*_XY_ as well as *P*_A_ and the degree of linkage between the *S* and *P* loci on both chromosomes (Figure 2A; Supplementary Figure 2). When *p*_*XY*2_ is strongly disfavored when maternally inherited (highly negative *a*_XY_) and/or strongly favored when paternally inherited (highly positive *b*_XY_, as determined by a higher *c*_XY_), then divergence at *P*_XY_ due to linkage to *S*_XY_ may protect *Y* from being replaced by *A*. However, if *p*_*A*2_ is sufficiently deleterious when maternally inherited (e.g. *a*_A_ ≫ *a*_XY_), then invasion of *A* may be favored. The scope for invasion of *A* is additionally increased when *p*_*A*2_ has stronger benefits when paternally inherited, i.e. for higher values of *c*_A_.

**Figure 2:**
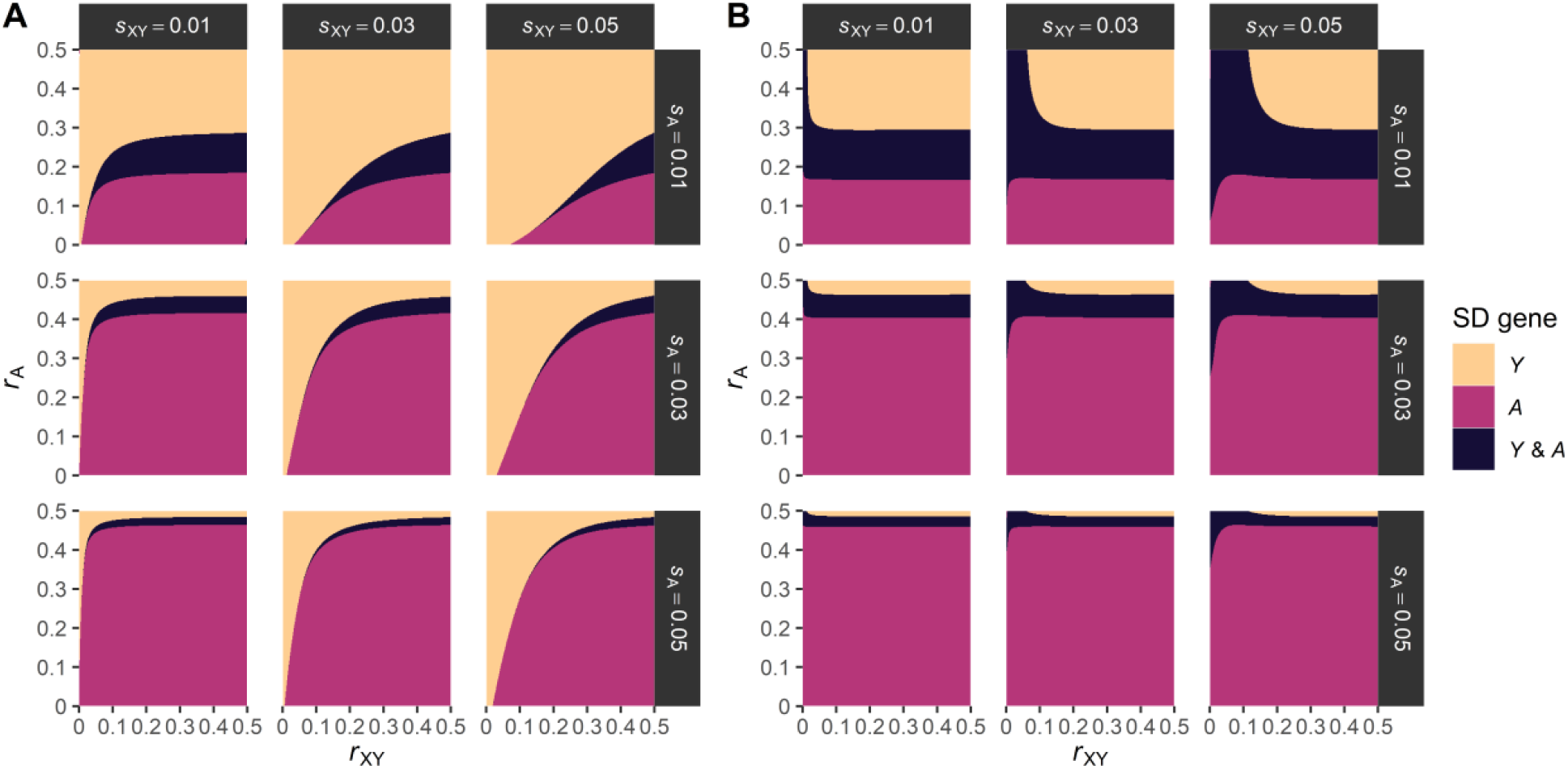
Invasion of a novel sex-determining allele *A* under different selection effect sizes and recombination rates. (A) Invasion of a male-determining *A*. (B) Invasion of a female-determining *A*. Different colors indicate whether *A* can invade and replace *Y* as the male-determining gene (pink) or not (orange), or whether a polymorphism occurs (dark blue). *Y/A* polymorphism is said to occur when both (A) *Y* and *A* have a frequency of at least 10% on the paternal allele of males or (B) *A* has a frequency between 10% and 90% on the maternal copy of females. Parameter values: *s*_XY_ = *s*_A_ = 0.02; *a*_XY_ = −0.1; *a*_A_ = −0.9; *c*_*XY*_ = *c*_*A*_ = 1.5. Fitted GAMs used *r*_XY_ and *r*_A_ as predictor variables; separate GAMs were fitted for each panel (i.e. combination of *s*_XY_ and *s*_A_).

Varying the selective effects *s*_*i*_, and the recombination rates *r*_*i*_ revealed that *A* can invade when linkage between *S*_A_ and *P*_A_ is sufficiently strong and/or *A* is associated with a selective effect *s*_A_ that sufficiently exceeds *s*_XY_ (Figure 2A). In contrast, stronger linkage between *S*_XY_ and *P*_XY_ or higher *s*_XY_ leads to decreased scope of invasion for *A*. Polymorphic sex determination systems, where *Y* and *A* coexist in the population, can also occur, particularly when linkage between *S*_XY_ and *P*_XY_ and between *S*_XY_ and *P*_XY_ is relatively low. The scope for *Y-A* polymorphism is however reduced when the selective effects associated with *P*_XY_ or *P*_A_ are higher.

### Transitions from male to female heterogamety

When *A* represents a female-determining gene that is dominant over *Y*, its invasion means that the ancestral male heterogamety system is replaced by female heterogamety system (Figures 1D, Supplementary Figure 2B), which additionally features fixation of *Y* in both sexes. Unlike the case when *A* represents a male-determining gene, I find that invasion of a female-determining *A* is virtually unconstrained (Figure 2B; Supplementary Figure 3) provided that recombination between *S*_A_ and *P*_A_ is sufficiently low. Only when recombination is high and selection on *P*_A_ is sufficiently weak can *Y* be maintained as the sex-determining gene.

One possible explanation is that the dominant feminizing *A* invades through two processes. Initially, divergence of *P*_*XY*_ has resulted in the formation of a coadapted *Y* − *p*_*XY*2_ haplotype that is restricted to males. Although *P*_*XY*_ remains polymorphic, meaning *p*_*XY*1_ and *p*_*XY*2_ are present in the population, the transmission of *p*_*XY*2_ from fathers to daughters will be reduced as *X p*_*XY*1_//*Y p*_*XY*2_ males have higher fitness than *X p*_*XY*2_//*Y p*_*XY*2_ males. This leads to a genetic load in females, which is resolved in females with a feminizing *A* as these may inherit the *Y* − *p*_*XY*2_ haplotype through the paternal route. This drives the initial invasion of *A*. At some point during its invasion, *A* becomes associated with *p*_*A*1_ which is beneficial when maternally inherited, driving *A* to fixation.

The invasion of *A* however induces a genetic load in males for similar reasons as applied to females under *XY* male heterogamety. One possible consequence is that a male-determining variant of *Y* that is dominant over the newly-invaded feminizing *A* should be able to invade. To test this, I determined whether (1) a dominant female-determining *A* could invade in a population with male heterogamety with *Y* as the male-determining gene, and (2) an even more dominant male-determiner *Y*^*\**^ could invade in a population where *A* had previously invaded as the female-determining gene. Such reciprocal invasions are indeed possible (Figure 3). Given that *A* can invade under an extremely broad range of parameter values when linkage is sufficiently strong (Figures 2B, Supplementary Figure 3), the scope for such dynamics are also likely to be broad.

**Figure 3:**
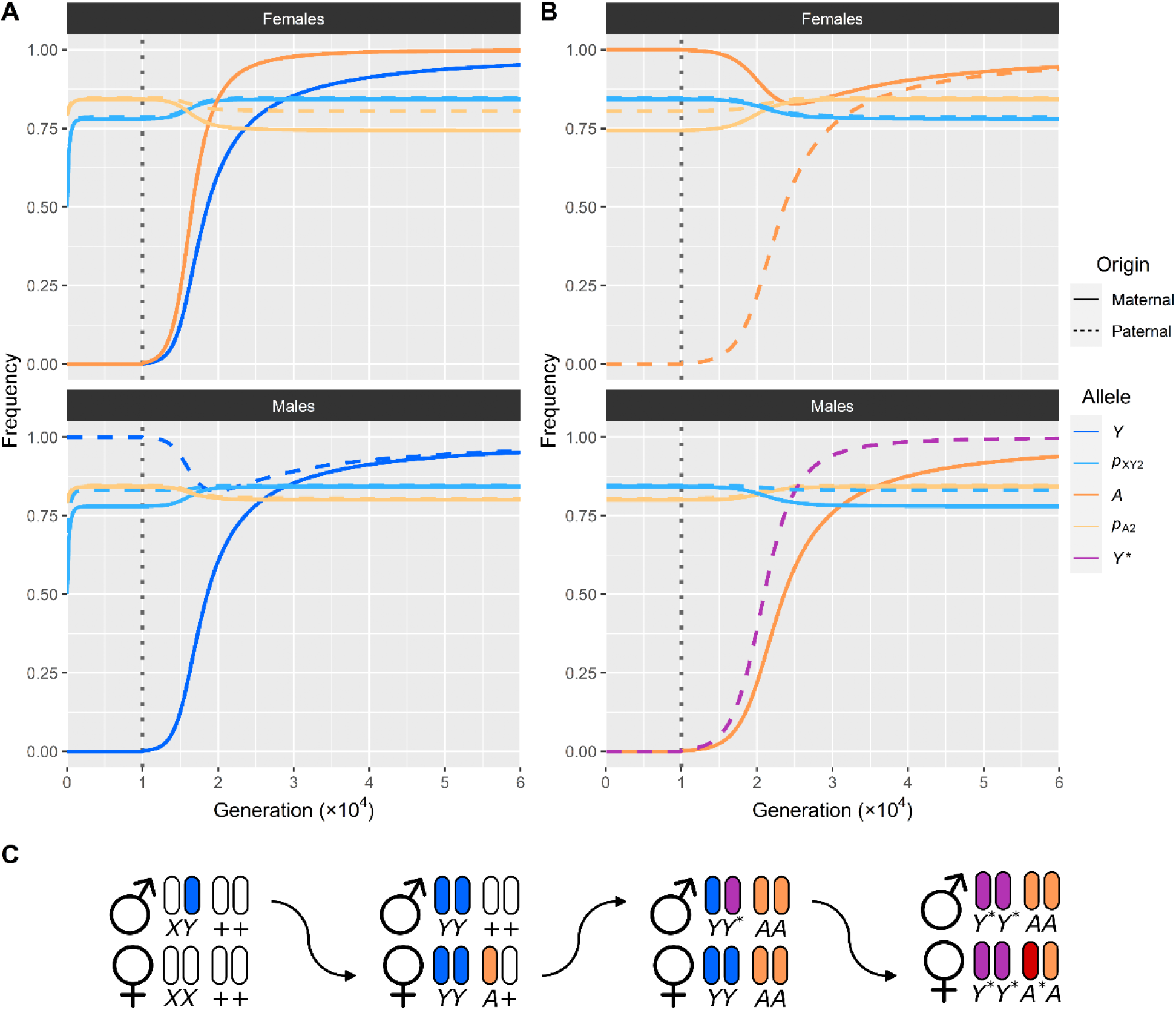
Sex chromosome ping pong through recurrent reciprocal turnover. (A) Invasion of a female-determining *A* that is dominant to *Y*. Fixation of Y is expected to take place over an extended period of time (not shown). (B) Invasion of a male-determining variant *Y\** that is dominant over the female-determining allele *A* that invaded in (A). The regular *Y* is presumed to be fixed prior to invasion of *Y\** (not shown). Dashed vertical lines denote introduction of *A* (in (A)) and *Y\** (in (B)). Parameter values: *s*_XY_ = *s*_A_ = 0.01; *r*_XY_ = *r*_A_ = 0.05; *a*_XY_ = *a*_A_ = −0.5; *c*_XY_ = *c*_A_ = 1.9. (C) Reciprocal invasibility leads to continuous alternations between *S*_XY_ and *S*_A_ as the most-dominant sex-determining gene. Arrows indicate transitions between male and female heterogamety and vice versa. From an initial population with *XY* male heterogamety (*Y* shown in blue), invasion of a feminizing *A* (orange) can occur which causes a transition from male to female heterogamety as *Y* is fixed in both sexes (similar to (A)). Subsequent invasion of a *Y\** (purple) that is dominant over *A* re-establishes male heterogamety (similar to (B)), after which a secondary feminizing *A\** (red) that is yet again dominant over *Y\** can lead again yield female heterogamety. Such patterns can in principle repeat indefinitely, establishing a ping pong pattern where the different chromosome pairs take turns as the sex chromosome pair.

## Discussion

Here, I presented a model to study transitions between sex determination mechanisms due to linkage to alleles with parentally antagonistic effects. Here, a novel sex determination gene is linked to a gene under parentally antagonistic selection. This proto-sex chromosome invades and replaces the pre-existing sex chromosome (which similarly carries a sex determination gene and a parentally antagonistic gene), establishing a novel sex chromosome system. Transitions between different chromosomes can occur through invasion of a sex determination gene with a homologous function, in which case the homogametic and heterogametic sex do not change (e.g. male heterogamety to male heterogamety), or through the invasion of a dominant gene that overrules the function of the ancestral sex determination gene, so that the homogametic and heterogametic sex switch (e.g. male heterogamety to female heterogamety).

Both types of sex determination turnovers can readily take place under parental antagonism, though the scope for invasion of a novel sex determination gene differs substantially. For a transition to a homologous sex determination gene, I find that invasion of the novel male-determining *A* can invade provided that the selective effects involved with *P*_A_ are sufficiently stronger than those of *P*_XY_, and/or the linkage to between *P*_A_ and *S*_A_ is tighter than that between *P*_XY_ and *S*_XY_ (Figure 2; Supplementary Figure 2). Transitions between different male heterogamety systems may be constrained, as invasion of the novel male-determining *A* requires that the co-adapted gene complex *Y* − *p*_XY2_ is broken down. That is, the sex-specific inheritance patterns of the XY chromosome pair promotes their differentiation. Here, the Y-chromosome acquires paternal-benefit alleles and the X-chromosome acquires maternal-benefit alleles. *A*-bearing males may lack the beneficial *Y* − *p*_XY2_ haplotype, instead carrying two complex *X* − *p*_XY1_ haplotypes at the XY chromosome pair. This leads to reduced fitness, and invasion of *A* is thus only favored if the initial benefit of inheriting an *A* − *p*_A2_ haplotype is sufficiently strong and/or reliable (i.e., unlikely to be broken down by recombination).

In contrast, transitions where the novel sex determination gene is dominant over the ancestral gene (and hence a change in heterogametic sex occurs) are feasible across virtually the entire parameter space considered here. One possible explanation (see also Results) is that the differentiation of the ancestral Y-chromosome leads to linkage disequilibrium between *Y* and the paternal-benefit allele *p*_XY2_. This establishes a co-adapted gene complex in males, particularly when paired with an X-chromosome with the maternal-benefit allele *p*_XY1_ as the *p*_XY1_*p*_XY2_ genotype has optimal fitness. Daughters from such males experience a genetic load, as paternal inheritance of *p*_XY1_ is disfavored. When a dominant feminizing allele *A* evolves, this genetic load can be resolved as the *Y* − *p*_XY2_ complex can now be transmitted to females. Such an effect was reported in pygmy mice, though the authors did not consider parental antagonism as an explanation for the benefit of Y-chromosomes in females (Saunders *et al*., 2014). This means that *A*-bearing females tend to have higher fitness than non-*A*-bearing females, promoting its initial spread in the population. As *A* persists, selection tends to favor those haplotypes where *A* is paired with *p*_XY1_ over those with *p*_XY2_. This differentiation can now occur, as *A* is always a female-limited gene. Consequently, *A* invades through the effect of two different selective processes.

One consequence of the spread of *A* is that males, rather than females, are now subject to a genetic load. Under these conditions they more often inherit the *p*_A2_ allele through their mothers, as *A p*_A1_//+ *p*_*a*2_ females have higher fitness than *A p*_A2_//+ *p*_*a*2_ females. These conditions enable the invasion of a new male-determining variant of *Y*, dubbed *Y*^*\**^, which is dominant over *A* (Figure 3). Effectively, this leads to dynamics where *S*_XY_ and *S*_A_ take turns as the dominant sex determination gene, as each invasion at one chromosome begets a new invasion at the other chromosome, with increasing levels of dominance of each newly-invading sex determination gene. This establishes a ‘sex chromosome ping pong’ where there are continuous switchovers between sex chromosome pairs and male versus female heterogamety (Figure 3C). This can lead to continuous evolution of both sex determination genes, and may help explain why some sex determination genes exhibit such high evolutionary rates, without invoking any conflict between them.

These effects are particularly interesting in light of the expected accumulation of parentally-antagonistic genes on sex chromosomes (Patten & Haig, 2009; Haig *et al*., 2014). As sex chromosomes develop from small sex-linked regions into genetically-distinct, non-recombining chromosomes, the genetic content of the X- and Y-chromosomes (or Z- and W-chromosomes in female heterogametic systems) is expected to diverge substantially. This could include the accumulation of genes with parentally-antagonistic fitness effects. If so, the divergence of these sex chromosomes does not render them more stable against turnover, but rather primes them for replacement.

Here, I considered that sex determination genes themselves did not affect fitness, but instead were linked to parentally antagonistic genes, which determined their ability to invade c.q. to resist invasion by other sex determination genes. From a functional perspective at least, sex determination genes may themselves exhibit parent-of-origin effects, as seen in several insect species where provision of sex determination gene transcripts is required for the successful execution of sex determination. For example, in the housefly *Musca domestica*, maternal provision of a female-specific *transformer* mRNA is required to ensure the autoregulatory feedback loop of *transformer* is properly initiated (Dübendorfer & Hediger, 1998; Hediger *et al*., 2010). In Hymenoptera, a sex determination mechanism known as maternal-effect genomic imprinting sex determination relies on the differential imprinting of a female-determining gene (Beukeboom *et al*., 2007). In these systems, the copy inherited from the mother is always inactivated, while the paternally inherited copy is active. Consequently, haploid unfertilized eggs carry an inactive feminizer and develop into males, whereas diploid fertilized eggs carry an active copy that is paternally inherited. Recently, such a gene has been identified in *Nasonia vitripennis* (Zou *et al*., 2020). An analysis of the potential evolutionary history of the sex determination mechanism in *Drosophila melanogaster* previously explored the role sex-specific selection may have played in stepwise successions of different sex determination genes and their variants (Pomiankowski *et al*., 2004). The findings here may similarly help explain why some sex determination genes evolve to exhibit parent-of-origin effects in terms of proximate function. Sex determination cascades were commonly thought to evolve bottom-up, with downstream components being conserved and upstream components being increasingly variable (Wilkins, 1995). However, downstream sex determination genes are not fully constrained in their evolution, and may still evolve novel functions even when under the control of other genes (Herpin *et al*., 2013; Schenkel *et al*., 2023). Potentially, some of the parent-of-origin effects on sex determination gene function reflect adaptations to parentally-antagonistic selection. While these effects were not formally integrated into my model, the finding that parentally-antagonistic selection drive transitions in sex determination makes it plausible that genes with parent-of-origin effects may have outcompeted variants that lacked such effects due to parentally-antagonistic selection.

Altogether, the results presented here contribute to our increasing understanding of the malleability of sex determination through numerous selective processes. In comparison to other models, sex determination transitions mediated by parental antagonism exhibit some very unusual dynamics, most striking of which is the possibility for different chromosome pairs to take turns as the sex chromosome pair. This can help explain why some sex determination cascades have genes that exhibit high evolutionary rates. As parental antagonism is only poorly understood, the prevalence of sex determination transitions that are in fact driven by this phenomenon is still unclear. However, as between-parent conflict is nearly ubiquitous, the scope for parental antagonism to occur may also be broad, and therefore parental antagonism may be a previously unconsidered factor in shaping sex determination mechanisms. As parental antagonism may act alongside other selective processes affecting sex determination genes, the peculiar dynamics described here may help understand why some sex chromosomes systems are so easily displaced.

## Acknowledgements

I acknowledge the financial support of the John Templeton Foundation (#62220). The opinions expressed in this paper are those of the author and not those of the John Templeton Foundation. I thank Femke Noorman for assistance with pilot studies on the model; the Center for Information Technology of the University of Groningen for providing access to the Hábrók high-performance computing cluster; and Manus Patten for useful comments on an earlier version of this manuscript.

## Data availability

Model source code, secondary data, analysis scripts, and output files are freely available through GitHub (https://github.com/MartijnSchenkel/SexDeterminationParentalAntagonism) and will be stored in Dryad upon acceptance.

## Supplementary Methods

### Model initialization

The model presented here is a modified version of that presented by Schenkel et al. (2021). It features two linkage groups, each of which consists of two loci, being the (potential) sex determination locus and the parentally antagonistic locus. All loci feature two possible alleles, which can be denoted using a 0 or 1. Each linkage group therefore features 4 potential haplotypes, each consisting of an allele at the SD locus and an allele at the parentally antagonistic locus. I denote non-sex determining alleles *+* on both linkage groups with a 0 and the sex-determining alleles *Y* and *A* with a 1. Similarly, the parentally antagonistic alleles *p*_*i*1_ and *p*_*i*2_ at linkage group *i* (1 = XY, 2 = A) are indicated using 0 and 1 respectively, so that e.g. a 11 haplotype consists of a dominant sex-determining allele (*Y, A*) and an allele *p*_*i*2_, and a 00 haplotype consists of the recessive (non-sex-determining) allele *+* and an allele *p*_*i*1_. I define an array ***G*** for which each element *g*_*ijk*_ gives the initial frequency of the *j*th haplotype (1 = 00, 2 = 01, 3 = 10, = 11) on linkage group *i* on the maternal (*k* = 1) and paternal (*k* = 2) copy. Using ***G***, I define our initial population ***P***, an array with dimension ***H***_mat_ × ***H***_pat_ in which ***H***_mat_ and ***H***_pat_ are 4 × 4 arrays wherein each element *h*_*ij*_ gives the frequency of a haplotype *i* on linkage group XY and *j* on linkage group A; subscripts mat and pat are used to distinguish between frequencies of the maternal and paternal copies of these haplotypes. Consequently, each element *p*_*ijkl*_ in ***P*** gives the frequency of the genotype that consists of haplotypes *i* and *k* for the maternal c.q. paternal copy of linkage group XY, and similarly *j* and *l* for linkage group A. Per locus *m* (1 = *S*_*XY*_, 2 = *P*_*XY*_, 3 = *S*_*A*_, 4 = *P*_*A*_), I define an array ***N***_*m*_ that counts the number of focal alleles (*Y, A, p*_*i*2_) in each element of ***P***, so that the inner product ***P*** · ***N***_*m*_ gives the population frequency of the focal allele at that locus. ***N***_*m*_ can be further split up into 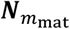 and 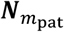, which give the number of the focal alleles on the maternal and paternal copies. Combinations (and where applicable transformations) of different ***N***_*m*_ arrays (and possibly the sex-determining arrays ***S***_M_, ***S***_F_; see details below) can be used to track the frequencies of different haplotypes in different sexes and of different parental origins (e.g., the entrywise product of 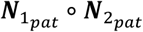 gives the frequency of a paternally-inherited haplotype with alleles *S*_*XY*_ and *p*_*XY*2_).

### Sex determination

Sex is determined by the number of focal alleles at the *S*_*XY*_ and *S*_*A*_ loci, and depends on whether A is a male-determining allele with identical function to *Y*, or a female-determining allele that is dominant to *Y*. I define a binary array ***S***_M_ which denotes whether a genotype *p* in ***P*** is male (1) or not (0); the binary array ***S***_F_ = 1 − ***S***_M_ indicates whether a genotype is female. If *A* has a male-determining function, then a genotype in ***P*** is male if ***N***_1_ > 0 ∨ ***N***_3_ > 0, and female otherwise. If *A* has a female-determining function, a genotype in ***P*** is male if ***N***_1_ > 0 ∧ ***N***_3_ = 0, and female otherwise. The entrywise products ***P***_F_ = ***P*** ∘ ***S***_F_ and ***P***_M_ = ***P*** ∘ ***S***_M_ represent the frequencies of genotypes among females (***P***_F_) and males (***P***_M_). Similar to the population-level frequency of the focal allele at locus *m*, the frequencies of the focal alleles among females and males are given by the inner products of ***P***_F_ · ***N***_*m*_ and ***P***_M_ · ***N***_*m*_.

### Fitness and selection

Fitness is determined by the genotypes at the *P*_XY_ and *P*_A_ loci. For each locus *i*, I define a vector *w*_*i*_ = {1, 1 + *a*_*i*_*s*_*i*_, 1 + *b*_*i*_*s*_*i*_, 1 + *s*_*i*_} that gives the fitness scores of respectively genotypes *p*_*i*1_ *p*_*i*1_, *p*_*i*2_*p*_*i*1_, *p*_*i*1_*p*_*i*2_, and *p*_*i*2_*p*_*i*2_ (where the initial allele indicates the maternal copy and the second allele the paternal copy). Consequently, *s*_*i*_ is the selective effect of the *p*_*i*2_ allele in homozygotes, and *a*_*i*_ and *b*_*i*_ are modifiers that determine the selective cost or benefit of the *p*_*i*2_ allele in heterozygotes when maternally or paternally inherited. I assume that *p*_*i*2_ has a fitness costs in heterozygotes when maternally inherited (*a*_*i*_ < 0), but a fitness benefit when paternally inherited (*b*_*i*_ > 1). An array ***W***_XY_ contains the fitness scores of each genotype in ***P*** based on the genotype at *P*_XY_, and similarly ***W***_A_ for the genotype at *P*_*A*_. These locus-specific fitness scores are assumed to be multiplicative, so that their entrywise product yields an array ***W*** = ***W***_XY_ ∘ ***W***_A_ that gives the total fitness for each genotype in ***P***. The entrywise product ***A***_F_ = ***W*** ∘ ***P***_F_ gives the frequency of each genotype among females after selection has taken place, and similarly ***A***_M_ = ***W*** ∘ ***P***_M_ for the genotype frequencies among adult males.

### Gametogenesis and reproduction

Reproduction occurs through random fusion of oocytes with sperm. Gametogenesis in males and females occurs in identical ways. To this end, I define an array ***U*** in which element *u*_*ijkl*_ that defines the probability of sampling a haplotype *l* from a genotype consisting of maternal haplotype *j* and paternal haplotype *k* on linkage group *i* (1 = XY, 2 = A), whilst accounting for recombination *r*_*i*_ between *S*_*i*_ and *P*_*i*_. Based on ***U***, I define an array ***T***_*ijklmn*_ = *u*_1*ikm*_ × *u*_2*jln*_. The matrix product of ***T*** with ***P***_F_ yields the frequency of gametes among oocytes, i.e. ***H***_F_ = ***TA***_F_, and similarly with males to obtain the gamete frequencies among sperm ***H***_M_ = ***TA***_M_. Note that ***H***_F_ and ***H***_M_ are functionally equivalent to ***H***_mat_ and ***H***_pat_, as both pairs represent the frequency of maternally and paternally inherited haplotypes. The Kronecker product ***H***_F_ ⊗ ***H***_M_ yields an array ***O*** that denotes the frequency of each genotype among the offspring. ***O*** has identical dimensions to ***P***, and effectively represents its offspring. Redefining ***P*** = ***O*** represents moving the simulation forward by 1 generation. All simulations are carried out for at least 50,000 generations.

I introduce the novel sex-determining allele *A* at generation 10,000 by manipulating the gamete arrays ***H***_F_ and ***H***_M_. For each, I redefine *h*_*i*3_ = *h*_*i*1_ × *μ* and *h*_*i*4_ = *h*_*i*2_ × *μ*, and subsequently *h*_*i*2_ = *h*_*i*2_ × (1 − *μ*) and *h*_*i*4_ = *h*_*i*4_ × (1 − *μ*) to convert a proportion *μ* of 00 and 01 (i.e., + *p*_21_ and + *p*_22_) gametes into 10 and 11 (*A p*_21_ and *A p*_22_).

## Supplementary Tables

**Supplementary Table 1:** Overview of model variables.

| **Supplementary Table 1:** Overview of model variables. |  |  |
| --- | --- | --- |
| **Parameter** | **Description** | **Range/standard value** |
| $i$ | Index describing the linkage group that a given variable refers to | {XY, A} |
| $S_i$ | Sex determination locus on linkage group $i$ | - |
| $P_i$ | Parentally antagonistic locus on linkage group $i$ | - |
| $r_i$ | Recombination between $S_i$ and $P_i$ on linkage group $i$ | [0,0.5] |
| $p_{i1}$ | Non-focal allele on linkage group $i$ | - |
| $p_{i2}$ | Focal allele with parentally antagonistic fitness effects on linkage group $i$ | - |
| $s_i$ | Selection coefficient for allele $p_{i1}$ | [0.01, 0.05] |
| $a_i$ | Dominance coefficient for $p_{i2}p_{i1}$ heterozygotes | [-1,0] |
| $b_i$ | Dominance coefficient for $p_{i1}p_{i2}$ heterozygotes | $-c_i a_i$ |
| $c_i$ | Scaling parameter used to calculate the value of $b_i$ based on $a_i$ | [1.1, 1.9] |
| $w_{i11}$ | Locus-specific fitness component of genotype $p_{i1}p_{i1}$ | 1 |
| $w_{i12}$ | Locus-specific fitness component of genotype $p_{i1}p_{i2}$ | $1 + b_i s_i$ |
| $w_{i21}$ | Locus-specific fitness component of genotype $p_{i2}p_{i1}$ | $1 + a_i s_i$ |
| $w_{i22}$ | Locus-specific fitness component of genotype $p_{i2}p_{i2}$ | $1 + s_i$ |

## Supplementary Figures

**Supplementary Figure 1:**
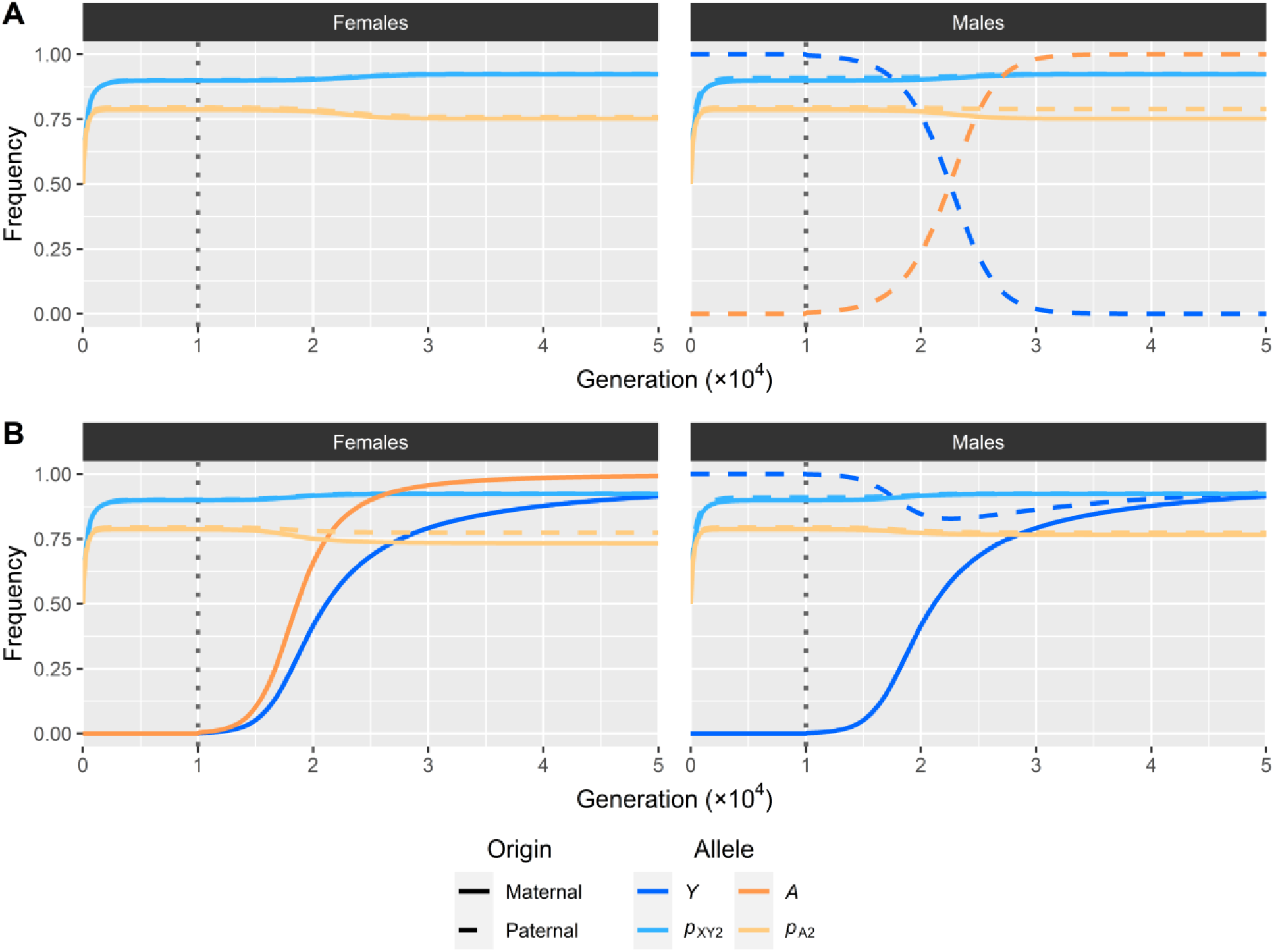
Dynamics of invasion by *A*. (A) Invasion of a male-determining allele *A* and a transition between homologous male heterogametic systems; *Y* is lost as *A* invades. (B) Invasion of a dominant female-determining allele *A* and a transition from male to female heterogametic system; *Y* approaches fixation as *A* invades. Note that the strength of selection for *Y* to increase in frequency drops once *A* approaches fixation, so that *Y* is not fully fixed yet at 50,000 generations (40,000 generations after *A* initially evolved). Only alleles with non-zero frequencies for a combination of haplotype and sex are shown. The dashed vertical line indicates the evolution of *A* through mutation in generation 10,000. Parameter values: *a*_*XY*_ = −0.2; *a*_*A*_ = −0.8; *c*_XY_ = *c*_A_ = 1.9; *s*_*XY*_ = *s*_*A*_ = 0.01; *r*_*XY*_ = *r*_*A*_ = 0.1.

**Supplementary Figure 2:**
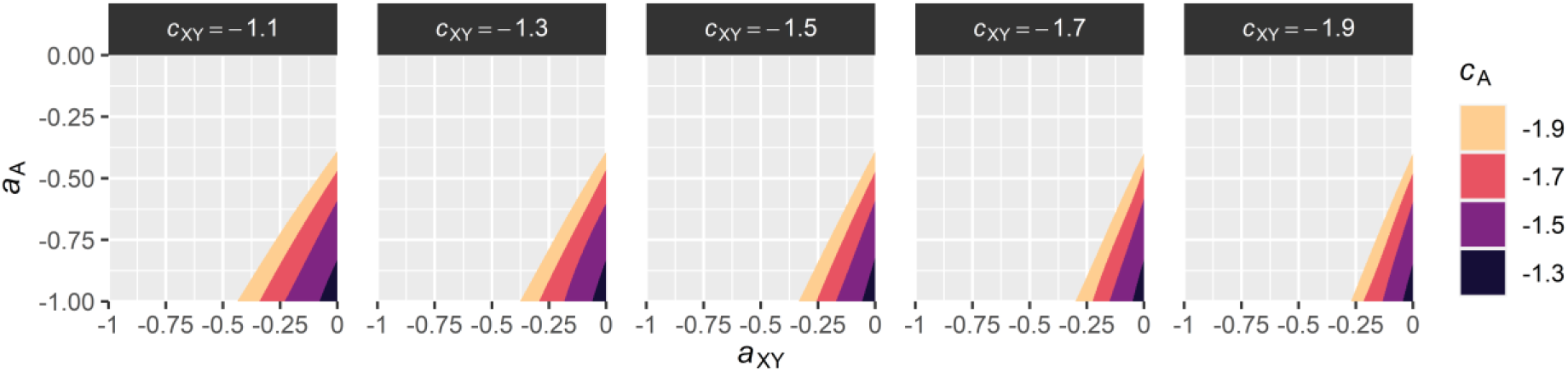
Scope for fixation of a male-determining allele *A* different parentally antagonistic selection regimes. Shaded areas represent the range of parameter values for which a male-determining *A* can invade. Parameter values: *s*_*XY*_ = *s*_*A*_ = 0.02; *r*_*XY*_ = *r*_*A*_ = 0.01. Fitted GAMs used *a*_XY_ and *a*_A_ as predictor variables; separate GAMs were fitted for each combination of *c*_XY_ and *c*_A_.

**Supplementary Figure 3:**
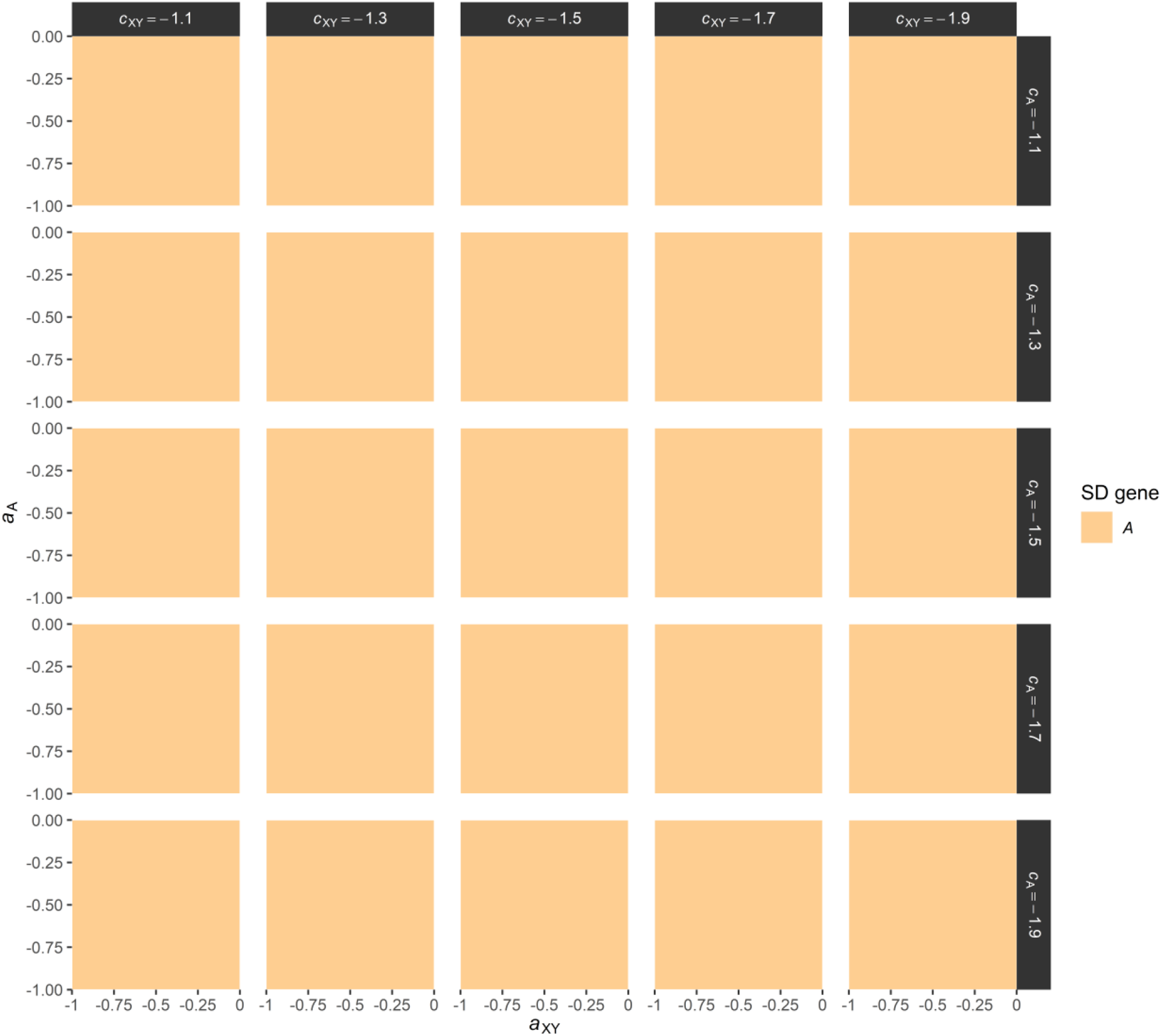
Scope for invasion of a female-determining allele *A*. For the entire parameter range considered here, *A* was able to invade and spread to fixation. Parameter values: *s*_*XY*_ = *s*_*A*_ = 0.02; *r*_*XY*_ = *r*_*A*_ = 0.01. Fitted GAMs used *a*_XY_ and *a*_A_ as predictor variables; separate GAMs were fitted for each panel (i.e. combination of *c*_XY_ and *c*_A_).

